# Decoding the brain state-dependent relationship between pupil dynamics and resting state fMRI signal fluctuation

**DOI:** 10.1101/2021.02.24.432768

**Authors:** Filip Sobczak, Patricia Pais-Roldán, Kengo Takahashi, Xin Yu

**Affiliations:** Translational Neuroimaging and Neural Control Group, High Field Magnetic Resonance Department, Max Planck Institute for Biological Cybernetics, 72076 Tuebingen, Germany; Graduate Training Centre of Neuroscience, International Max Planck Research School, University of Tuebingen, 72074 Tuebingen, Germany; Institute of Neuroscience and Medicine 4, Medical Imaging Physics, Forschungszentrum Jülich, 52425 Jülich, Germany; Athinoula A. Martinos Center for Biomedical Imaging, Massachusetts General Hospital and Harvard Medical School, Charlestown, MA 02129, USA

**Keywords:** pupil, fMRI, decoding, principal component analysis, neuromodulation, brain state

## Abstract

Pupil dynamics serve as a physiological indicator of cognitive processes and arousal states of the brain across a diverse range of behavioral experiments. Pupil diameter changes reflect brain state fluctuations driven by neuromodulatory systems. Resting state fMRI (rs-fMRI) has been used to identify global patterns of neuronal correlation with pupil diameter changes, however, the linkage between distinct brain state-dependent activation patterns of neuromodulatory nuclei with pupil dynamics remains to be explored. Here, we identified four clusters of trials with unique activity patterns related to pupil diameter changes in anesthetized rat brains. Going beyond the typical rs-fMRI correlation analysis with pupil dynamics, we decomposed spatiotemporal patterns of rs-fMRI with principal components analysis (PCA) and characterized the cluster-specific pupil-fMRI relationships by optimizing the PCA component weighting via decoding methods. This work shows that pupil dynamics are tightly coupled with different neuromodulatory centers in different trials, presenting a novel PCA-based decoding method to study the brain state-dependent pupil-fMRI relationship.

## Introduction

Pupil diameter changes reflect the brain state and cognitive processing (1–4). It contains information about behavioral variables as diverse as a subject’s arousal fluctuation (5–7), sensory task performance (5, 8), movement (9–12), exerted mental effort (13–15), expected reward (16), task-related uncertainty (17–19), or upcoming decisions (20, 21). This richness of behavioral correlates is partly explained by the fact that multiple neuronal sources drive pupil activity. Pupil diameter changes reflect spontaneous neural activity across the cortex (9–11, 22, 23) and in major subcortical areas (9, 24–27). Both sympathetic and parasympathetic systems innervate muscles controlling pupil dilation and constriction (28–30), and the activity of subcortical nuclei mediating neuromodulation has been tightly coupled with pupillary movements (23, 24, 31–34). In particular, rapid and sustained pupil size changes are associated with cortical noradrenergic and cholinergic projections respectively (31) and direct recordings of the noradrenergic locus coeruleus demonstrate neuronal activity highly correlated with pupil dynamics (24, 32, 35). Also, pupil diameter changes are regulated through dopaminergic neuromodulation under drug administration (36) and in reward-related tasks (16, 33). Studies also show that pupil dilation and constriction can be controlled by serotonergic agonists and antagonists, respectively (37, 38). These studies have revealed the highly complex relationship between pupil dynamics and brain state fluctuations (7, 12, 30, 39).

Resting state fMRI (rs-fMRI) studies have uncovered global pupil-fMRI correlation patterns in human brains as well as revealed that the pupil dynamics-fMRI relationship changed under different lighting conditions or when subjects engaged in mental imagery (22, 26). The dynamic functional connectivity changes detected by fMRI, possibly modulated by the interplay of cholinergic and noradrenergic systems (40), are also reflected by pupil dynamics both at rest (41) and in task conditions (42). Furthermore, rs-fMRI has been used to display a differential correlation pattern with brainstem noradrenergic nuclei, e.g., A5 cell group, depending on the cortical cross-frequency coupling state in the animal model (23). Although rs-fMRI enables brain-wide pupil-fMRI correlation analysis across different states, the linkage of brain state-dependent pupil dynamics with distinct activation patterns of neuromodulatory nuclei remains to be thoroughly investigated beyond the conventional analysis methods.

Here, we aimed to differentiate brain states with varied coupling patterns of pupil dynamics with the subcortical activity of major neuromodulatory nuclei in an anesthetized rat model. First, we demonstrated that the pupil-fMRI relationship is not uniform across different scanning trials and employed a clustering procedure to identify distinct pupil-fMRI spatial correlation patterns from a cohort of datasets. Next, we modeled the relationship of the two modalities for each cluster using principal component analysis (PCA)-based decoding methods (gated recurrent unit (GRU) (43) neural networks and linear regression) and characterized unique subcortical activation patterns coupled with specific pupil dynamic features. This work demonstrates the effectiveness of PCA-based decoding to dissect the time-varied pupil-fMRI relationship corresponding to different forms of brain state-dependent neuromodulation.

## Results

### Identification of brain states with distinct pupil dynamics correlation patterns

To investigate brain state-dependent pupil dynamics, we acquired whole-brain rs-fMRI with real-time pupillometry in anesthetized rats (n=10) as previously reported (23). Initially, the pupil dilation and fMRI time-series from all trials (n=74) were concatenated. A voxel-wise correlation map of the concatenated pupil dynamics signals with fMRI time courses showed a global negative correlation (Fig. 1A). However, the generated map was not representative of all trials, which was revealed by creating correlation maps for individual trials (Fig. 1B). These maps demonstrated high variability of pupil-fMRI correlations, which is presented by the histogram distribution of spatial correlation values between individual-trial spatial maps and the concatenated all-trial map (Fig. 1C).

**Fig. 1.**
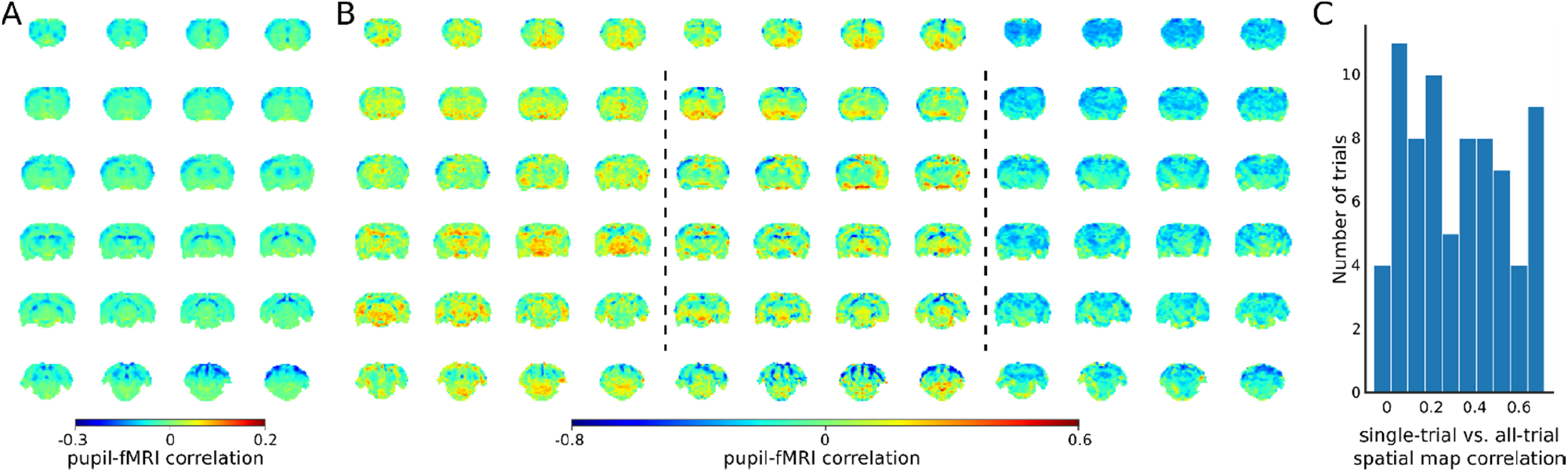
Variability of the pupil-fMRI linkage. (A) The pupil-fMRI correlation map created by correlating the two modalities’ concatenated signals from all trials. (B) Selected individual trial correlations maps. (C) Histogram of spatial correlations between the all-trial correlation map and individual trial maps. High variability of similarities between the maps shows that the pupil-fMRI relationship is not stationary and changes across trials.

Next, we clustered all trials into different groups based on pupil-fMRI correlation maps (Fig. 2A). To facilitate the clustering analysis we reduced the dimensionality of the spatial correlation maps using the uniform manifold approximation and projection (UMAP) (44) method and decreased the number of features used for clustering from the number of voxels (n=20804) to 72 for each map. Three to seven clusters were identified with Gaussian mixture modeling and examined using silhouette analysis (45, 46). Here, we focused on the 4-cluster categorization since this division yielded the highest mean silhouette scores (Fig. 2B). The power spectral density (PSD) estimates of pupil dynamics were plotted based on the cluster division in Fig. 2C. PSD of cluster 1 showed a distinct peak at 0.018 Hz as well as the lowest baseline pupil diameter values. In contrast, cluster 4 had the highest mean baseline diameter and a peak at 0.011 Hz. Clusters 2 and 3 showed peaks of oscillatory power at less than 0.01 Hz. The ultra-slow oscillation is typical for spontaneous pupil fluctuations (47). We recreated pupil-fMRI correlation maps based on the 4 clusters (Fig. 2D). Three clusters (1, 2 and 4) showed negative correlations across large parts of the brain, with the correlation strength differing across clusters. In contrast, cluster 3 displayed a very low mean correlation with positive coefficients spreading across the entire brain. It is also noteworthy that cluster 1 showed a high positive correlation in the periaqueductal gray and ventral midbrain regions. These results verified the usage of data-driven clustering for identifying brain state-dependent pupil dynamics.

**Fig. 2.**
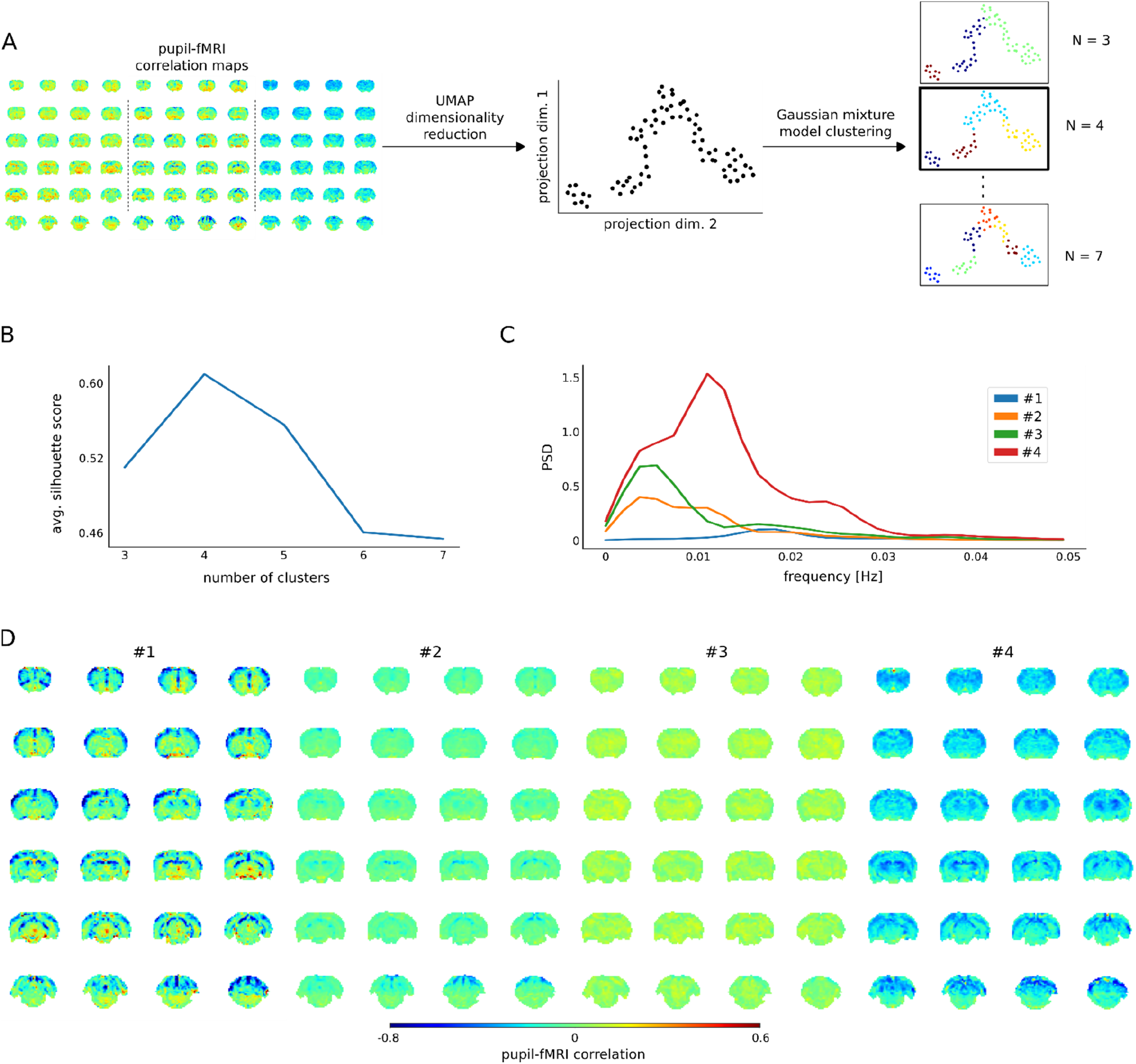
Clustering of trials with distinct pupil-fMRI correlation patterns. (A) Schematic of the clustering procedure. UMAP is used to reduce the dimensionality of all individual-trial correlation maps to 72 dimensions. A 2D UMAP-projection of the real data is shown. Each dot represents a single trial. The trials are clustered using Gaussian mixture model clustering. Different numbers of clusters are evaluated. (B) The final number of clusters is selected based on silhouette analysis. The highest average silhouette score is obtained by k=4 clusters. (C) Pupil power spectral density estimates (PSD) of each of the four clusters. Signals were downsampled to match the fMRI sampling rate. (D) Cluster-specific correlation maps based on concatenated signals belonging to the respective groups.

### Decoding-based investigation of the relationship between whole-brain rs-fMRI and pupil dynamics

To characterize the pupil-fMRI relationship beyond the conventional correlation analysis, we implemented data-driven decoding models to dissect the dynamics of the two modalities. First, we performed principal component analysis (PCA) to extract spatiotemporal features of whole-brain rs-fMRI signals (n=300) and trained either linear regression (LR) or a gated recurrent unit (GRU) neural network to predict pupil dynamics based on rs-fMRI PCA time courses (Fig. 3A). Furthermore, we compared the LR and GRU prediction models with a correlation template-based pupil dynamics estimation used in previous studies (23, 48). All methods were trained on 64 trials using cross-validation and then were tested on additional 10 independent trials. As the correlation template-based predictions were bounded to the <−1; 1> range, Pearson’s correlation coefficient was used to evaluate the decoding of all methods. We optimized the hyperparameters of GRUs and linear regression variants using Bayesian optimization (49, 50) and 4-fold cross-validation (hyperparameter values are listed in Methods). Both linear regression and GRU outperformed the correlation-template approach on both training (CC_base_=0.37±0.27 s.d., CC_LR_=0.45±0.26 s.d., CC_GRU_=0.46±0.25 s.d., p_LR_=4.3*10^−6^, p_GRU_=2.4*10^−6^) and test sets (CC_base_=0.25±0.17 s.d., CC_LR_=0.44±0.24 s.d., CC_GRU_=0.45±0.27 s.d., p_LR_=0.003, p_GRU_=0.01) (Fig. 3B). We also verified the number of rs-fMRI PCA components by testing varied component counts, showing that the highest prediction scores were achieved with 300 components (Supplementary Fig. 1). In addition, when varying the temporal shift between pupil dynamics and rs-fMRI signals, we obtained the highest prediction scores with zero shift between the input and output signals (Supplementary Fig. 1). Interestingly, the component which explained the most pupillary variance (explained var. = 7.03 %) and had the highest linear regression weight, explained only 0.8 % of the fMRI variance (Supplementary Fig. 2). Furthermore, the component which explained the most fMRI variance (explained var. = 22.01 %), was weakly coupled with the pupil fluctuation (explained var. = 0.51 %). Thus, this prediction-based PCA component weighting scheme enabled the dissection of unique brain activity features for the modeling of the pupil-fMRI relationship. It should also be noted that GRU and linear regression methods obtained comparable scores and both methods showed similar prediction performance (Fig. 3C). Supplementary Fig. 3 shows prediction maps created by integrating PCA components using linear regression weights or averaged GRU gradients (details in Methods). The resemblance of the two maps suggests that despite GRU’s potential for encoding complex and non-linear functions, a linear regression-based rs-fMRI mapping scheme was sufficient for predicting pupil dynamics.

**Fig. 3.**
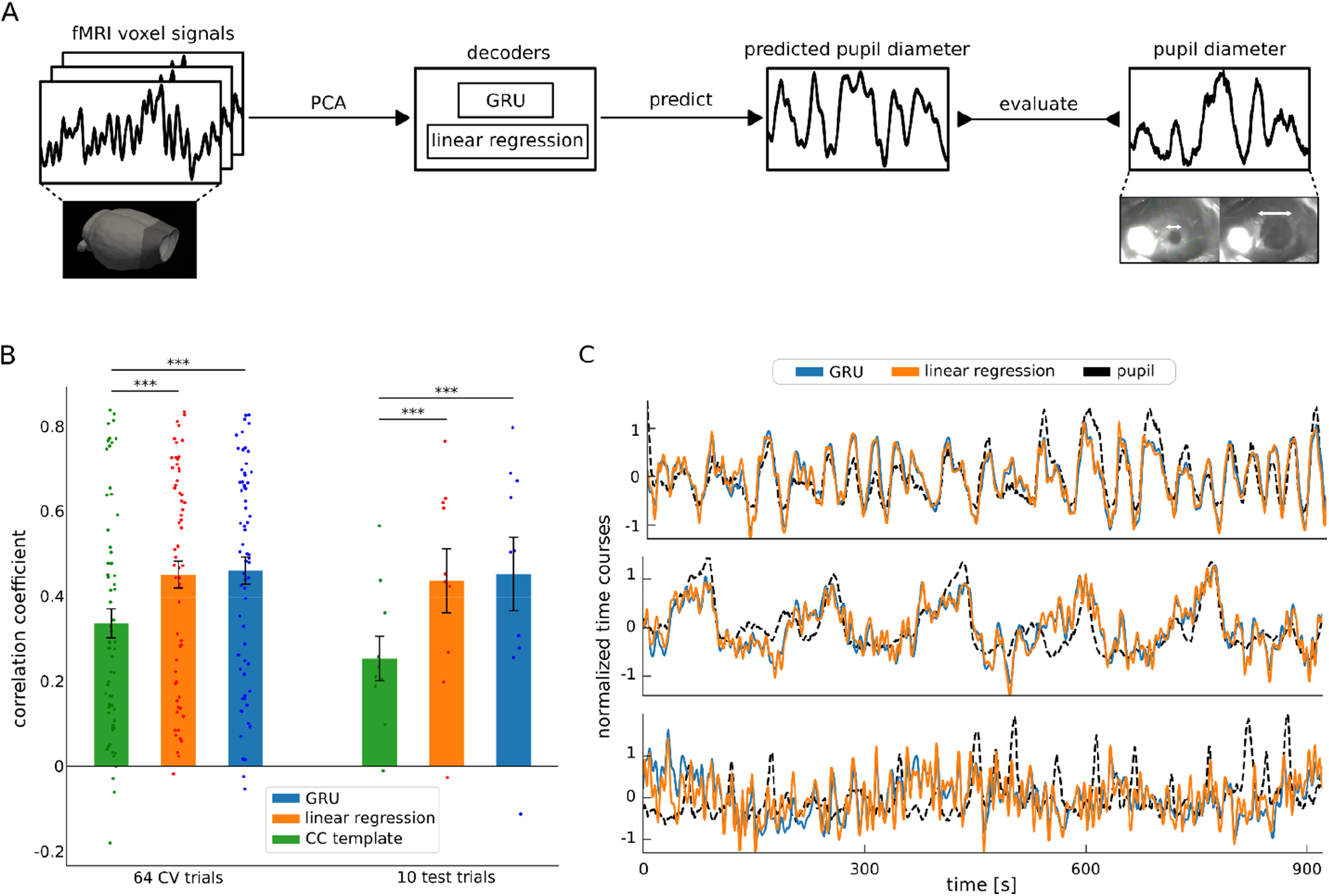
Decoding pupil dynamics based on fMRI signals. (A) Schematic of the decoding procedure. Principal component analysis (PCA) was applied to fMRI data from all trials. The PCA time courses were fed into either linear regression or GRU decoders which generated pupil signal predictions. The prediction quality was evaluated by comparing the generated signals with real pupil fluctuations using Pearson’s correlation coefficients. (B) Comparison of the three methods’ pupil dynamics predictions. Linear regression and GRU performed better than the correlation-based baseline method on both the cross-validation splits (CC_base_=0.37±0.27 s.d., CC_LR_=0.45±0.26 s.d., CC_GRU_=0.46±0.25 s.d., p_LR_=4.3*10^−6^, p_GRU_=2.4*10^−6^) and on test data (CC_base_=0.25±0.17 s.d., CC_LR_=0.44±0.24 s.d., CC_GRU_=0.45±0.27 s.d., p_LR_=0.003, p_GRU=0.01_). Scattered points show individual prediction scores. (C) Linear regression and GRU predictions of three selected trials (CC_GRU-top=0.79_, CC_LR-top=0.77_, CC_GRU-middle=0.75_,CC_LR-middle=0.73_, CC_GRU-bottom=0.02_, CC_LR-bottom=0.06_). Qualitatively, linear regression and GRU predictions are very similar.

The map generated by combining PCA components with the linear regression decoder allowed the identification of brain nuclei which were not highlighted in the correlation map shown in Fig. 1A. Fig. 4 shows an overview of the PCA-based fMRI prediction map overlaid on the brain atlas, revealing pupil-related activation patterns at key neuromodulatory nuclei of the ascending reticular activating system (ARAS) - the dopaminergic ventral tegmental area, substantia nigra and supramammillary nucleus, the serotonergic raphe and B9 cells, the histaminergic tuberomammillary nucleus, the cholinergic laterodorsal tegmental and pontine nuclei, the glutamatergic parabrachial nuclei and the noradrenergic locus coeruleus. Positive weights were also located in subcortical regions involved in autonomous regulation - the lateral and preoptic hypothalamus and the periaqueductal gray. In addition, the subcortical basal forebrain nuclei (the horizontal limb of the diagonal band, nucleus accumbens, and olfactory tubercle) and the septal area were positively coupled to pupil dynamics. Lastly, regions of the hippocampal formation - the hippocampus, entorhinal cortex and subiculum, as well as cingulate, retrosplenial and visual cortices displayed positive weighting. It should be noted that the thalamus and the hippocampus displayed both positive and negative weights. Negative coupling was also found in the cerebellum and most somatosensory cortical regions. The voxel-wise statistical significance (p<0.01) was validated using randomization tests and corrected for multiple comparisons with false discovery rate correction (details in Methods). These results highlight the advantage of using PCA decomposition combined with prediction-based decoding methods instead of conventional correlation analysis to identify pupil-related subcortical activation patterns.

**Fig. 4.**
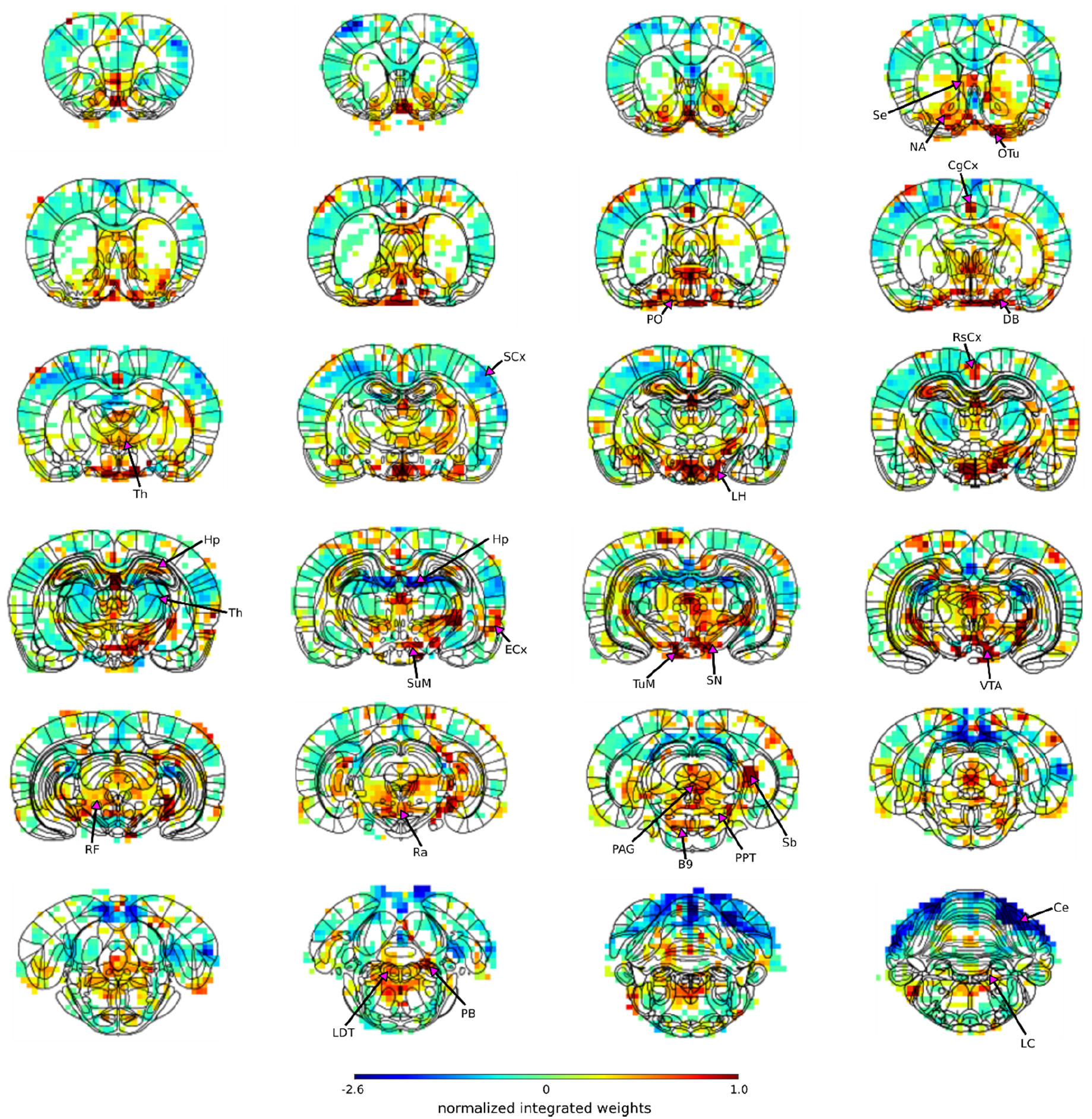
Localization of pupil dynamics-related information content across the brain. The spatial map highlights regions from which pupil-related information was decoded. It was created by integrating PCA spatial maps with weights of the trained linear regression model. The map displays positive weights in all positive neuromodulatory regions of the ascending reticular activating system as well as in other regions involved in autonomous regulation - the lateral and preoptic hypothalamus and the periaqueductal gray. The subcortical basal forebrain nuclei (the horizontal limb of the diagonal band, nucleus accumbens, and olfactory tubercle) and the septal area were also positively coupled to pupil dynamics. Finally, regions of the hippocampal formation - the hippocampus, entorhinal cortex and subiculum, as well as cingulate, retrosplenial and visual cortices showed positive weights. The thalamus and the hippocampus had both positive and negative weights. Strong negative weighting was found in the cerebellum and most somatosensory cortical regions. Masked regions (white) did not pass the false discovery rate corrected significance threshold (p=0.01). Abbreviations: B9 – B9 serotonergic cells, Ce – cerebellum, CgCx – cingulate cortex, DB – horizontal limb of the diagonal band, ECx – entorhinal cortex, Hp – hippocampus, LC – locus coeruleus, LDT – laterodorsal tegmental nuclei, LH – lateral hypothalamus, NA – nucleus accumbens, OTu – olfactory tubercle, PAG – periaqueductal gray, PB – parabrachial nuclei, PO – preoptic nuclei, PPT-pedunculopontine tegmental nuclei, Ra – raphe, RF – reticular formation, RsCx – retrosplenial cortex, Sb – subiculum, SCx – somatosensory cortex, Se – septal nuclei, SG – subgeniculate nucleus, SN – substantia nigra, SuM – supramammillary nucleus, Th – thalamus, TuM – tuberomammillary nucleus, VTA – ventral tegmental area.

### Characterization of brain state-dependent PCA-based pupil-fMRI prediction maps

To differentiate brain state-dependent subcortical activation patterns related to different pupil dynamics, we retrained the linear regression model based on the 4 different clusters shown in Fig. 2D and created PCA-based fMRI prediction maps for each cluster (Fig. 5).

**Fig. 5.**
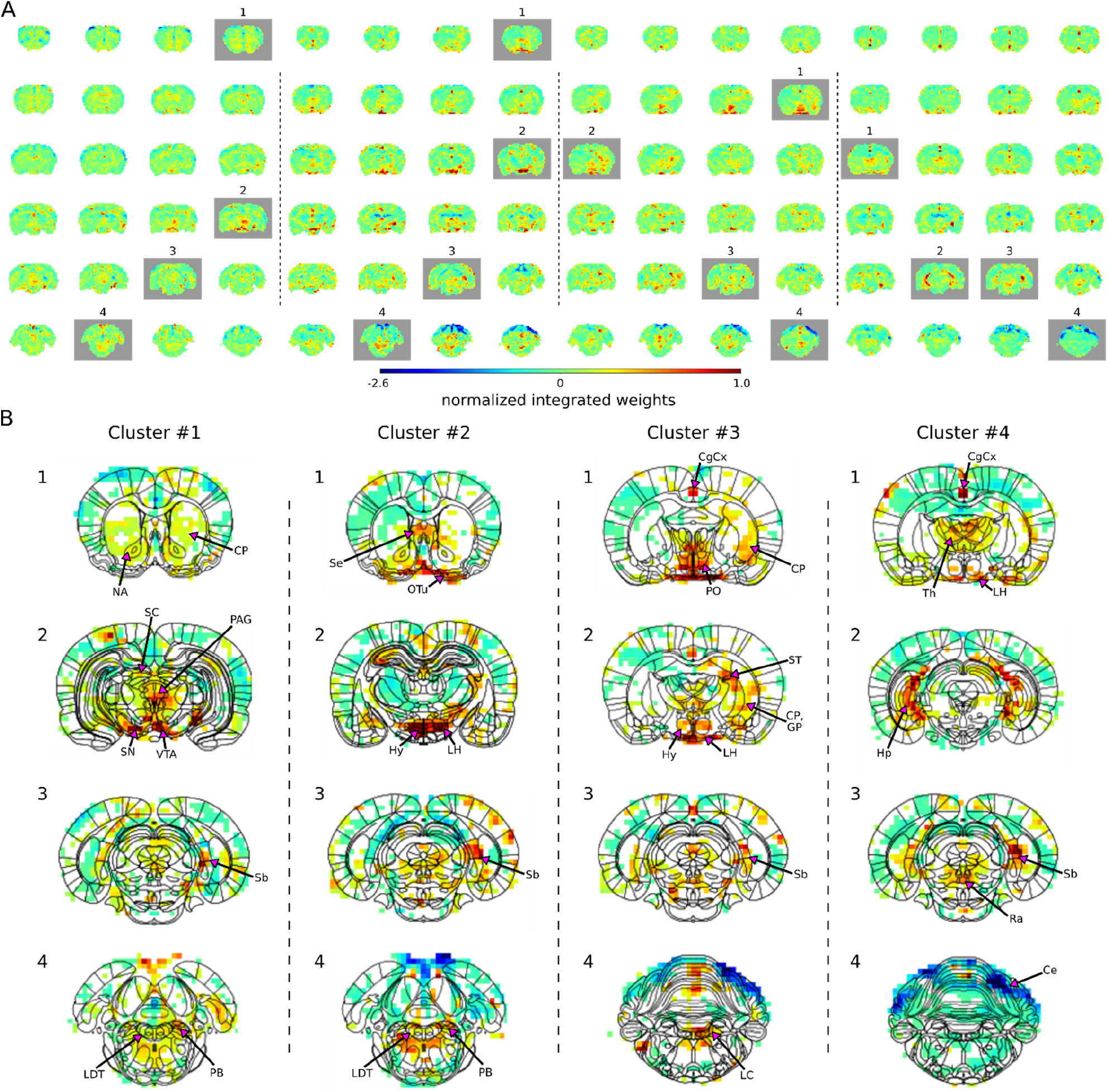
Characterization of brain state-specific pupil-fMRI relationships. (A) Pupil information content maps generated by integrating PCA spatial maps with weights of linear regression models trained on cluster-specific trials. In all clusters, negative weights were found in the somatosensory cortex, the cerebellum and posterior parts of the thalamus. Common to all clusters, were positive weights in anterior thalamic, preoptic and hypothalamic nuclei, the subiculum and parts of the hippocampus. Clusters 1-3 displayed positive weights in neuromodulatory brainstem regions, substantia nigra and ventral tegmental area, as well as the entorhinal cortex. The cingulate cortex and retrosplenial cortex were positive in clusters 2-4. Marked with gray are frames plotted in B. (B) Cluster-specific spatial patterns are portrayed on slices selected from A (marked with gray rectangles). Characteristic to cluster 1 were positive weights in the dopaminergic substantia nigra and ventral tegmental area as well as in their efferent projections in the nucleus accumbens and caudate-putamen. Positive weighting was also found in the periaqueductal gray and brainstem laterodorsal tegmental and parabrachial nuclei, as well as in the superior colliculus. Cluster 2 was characterized by the strongest positive weights in hypothalamic regions, lateral in particular. Brainstem arousal-regulating locus coeruleus, laterodorsal tegmental and parabrachial nuclei, as well as the septal area and the olfactory tubercle displayed high positive weights. In cluster 3, as in cluster 2, the locus coeruleus, laterodorsal tegmental and parabrachial nuclei showed positive linkage with pupil dynamics. The highest cluster 3 values were located in preoptic and other hypothalamic areas, as well as in stria terminalis carrying primarily afferent hypothalamic fibers, caudate-putamen and globus pallidus. In cluster 4 the only neuromodulatory region showing positive weights was the caudal raphe. The anterior parts of the brainstem displayed negative weighting. Characteristic to cluster 4 were high weights in the thalamus and in the hippocampus and the subiculum forming the hippocampal formation. Masked regions (white) did not pass the false discovery rate corrected significance threshold (p=0.01). Abbreviations: CgCx – cingulate cortex, CP – caudate-putamen, GP – globus pallidus, Hp – hippocampus, Hy – hypothalamus, LH – lateral hypothalamus, NA – nucleus accumbens, OTu – olfactory tubercle, PAG – periaqueductal gray, PO – preoptic nuclei, TSb – subiculum, SC – superior colliculus, Se – septal area, SN – substantia nigra, ST – stria terminalis, Th – thalamus, VTA – ventral tegmental area.

Each PCA-based prediction map portrayed a cluster-specific spatial pattern (Fig. 5B). Cluster 1 was characterized by strong positive weights in the dopaminergic substantia nigra and ventral tegmental area as well as in their efferent projections in the striatum (nucleus accumbens and caudate-putamen) (51). Positive coupling was also displayed in the periaqueductal gray and brainstem laterodorsal tegmental and parabrachial nuclei as well as in the superior colliculus. Cluster 2 had the strongest positive weights in hypothalamic regions, lateral in particular, but also in brainstem arousal-regulating locus coeruleus, laterodorsal tegmental and parabrachial nuclei. High positive values were also found in the septal area and the olfactory tubercle. In cluster 3 the highest values were visible in preoptic and other hypothalamic areas, as well as in stria terminalis carrying primarily afferent hypothalamic fibers (52), caudate-putamen and globus pallidus. As in cluster 2, the locus coeruleus, laterodorsal tegmental and parabrachial nuclei showed positive linkage with pupil dynamics. Contrastingly, in cluster 4 caudal raphe was the only neuromodulatory region showing positive weights and the anterior parts of the brainstem displayed negative weighting. Characteristic to cluster 4 were high weights in the hippocampus and the subiculum forming the hippocampal formation, as well as in thalamic and amygdaloid areas. Common to all clusters, negative weights were detected across somatosensory cortices, the cerebellum and posterior parts of the thalamus, as well as positive weights in hypothalamic and anterior thalamic nuclei. The subiculum and parts of the hippocampus were also positive in all clusters, however, the entorhinal cortex, also belonging to the hippocampal formation, was positive only in clusters 1-3. The same three clusters showed major positive weights in the neuromodulatory brainstem regions, substantia nigra, and ventral tegmental area. Clusters 2-4 displayed strong weights in the retrosplenial cortex and the cingulate cortex which has been coupled with both noradrenergic modulation (35) and pupil dynamics (23, 24). Here, we demonstrated the effectiveness of the PCA-based approach to reveal brain state-specific subcortical activity patterns related to pupil diameter changes.

## Discussion

Previous studies analyzed the relationship of fMRI and pupil dynamics either by directly correlating pupil size changes with the fMRI signal fluctuation (22, 23, 26) or by applying a general linear model to produce voxel-wise activation maps (15, 34, 53). Here, we performed PCA-based dimensionality reduction to decouple spatiotemporal features of fMRI signals (54) and implemented prediction methods to decode pupil dynamics based on the optimized PCA component weighting (Fig. 3).

Two advantages can be highlighted in the present pupil-fMRI dynamic mapping scheme. First, conventional correlation analysis relies on the temporal features of fMRI time courses from individual voxels or regions of interest. Hence, it could not decouple the superimposed effects of multiple signal sources (55, 56) or characterize the state-dependent dynamic subcortical correlation patterns. On the other hand, the PCA decomposition scheme solved these issues by decoupling multiple components of rs-fMRI signals with unique spatiotemporal patterns carrying pupil-related information. Second, the data-driven training of prediction methods optimized the weighting of individual rs-fMRI PCA components. Using the optimized neural network (GRU) or linear regression (LR)-based decoding models, we created prediction maps linking pupil dynamics with fMRI signal fluctuation of specific subcortical nuclei (Fig. 4, Supplementary Fig. 3). Also, the decoding models showed much better pupil dynamics prediction than the correlation template-based approach reported previously (23, 48). Meanwhile, it should be noted that both LR and GRU models generated qualitatively similar prediction maps, highlighting the pupil-related rs-fMRI signal fluctuation from the same subcortical brain regions (Fig. 3C, Supplementary Fig. 3). Unlike our previous single-vessel fMRI prediction study (57), the GRU-based neural network prediction scheme may require much bigger training datasets to outperform linear regression modeling (58). Another plausible explanation is that the pupil dynamics were predominantly and linearly driven by only a few rs-fMRI PCA components (Supplementary Fig 3), presenting brain activation patterns related to arousal fluctuation and autonomous regulation (59, 60).

The PCA-based prediction modeling provides a novel scheme to decipher subcortical spatial patterns of fMRI signal fluctuation related to brain state-dependent pupil dynamics. Most notably, neuromodulatory nuclei of ARAS and other subcortical nuclei involved in brain state modulation, as well as autonomous regulation were identified in the PCA-prediction map created from all trials. The highlighted hypothalamus, basal forebrain, and neuromodulatory brainstem nuclei are responsible for both global brain state modulation as well as autonomous cardiovascular, respiratory and baroreflex control (60–65). Consequently, the source of pupil-related information found across the cortex was probably modulated through global subcortical projections rather than a more direct causal interaction with pupil size changes (31, 66). The observed activation pattern suggests that, in the anesthetized state, pupil diameter fluctuation reflects a complex interaction of subcortical homeostatic and neuromodulatory centers.

Also, we have shown that these subcortical interactions and the neural correlates of pupil dynamics are not stationary but change across trials in a brain state-dependent manner. Based on the correlation patterns, we identified four clusters of trials with distinct pupil-fMRI coupling. Our results demonstrate that pupil size changes can be modulated by different combinations of subcortical nuclei, indicating varied brain state fluctuation underlying different oscillatory patterns of pupil dynamics (Fig. 2C). This is further exemplified by examining the cluster-specific PCA prediction maps. The map of cluster 2 demonstrates the strongest coupling of pupil dynamics with the hypothalamus, which is known to drive pupil dilation (27) and also highlights other brain state-regulating nuclei of the ARAS. It is possible that the hypothalamus was the key driver of brain state fluctuation in cluster 2 (65, 67). On the other hand, hypothalamic weights were least prevalent in cluster 1 which displayed strong pupil coupling with the dopaminergic system known to modulate pupil dynamics (16, 33, 36). Finally, in trials of cluster 4, the caudal raphe nucleus was the only brainstem neuromodulatory nucleus whose activity was positively weighted to predict pupil fluctuations. Additionally, the subiculum weights were the strongest in cluster 4 out of all clusters. The positive coupling of the raphe and subiculum hints at the possibility of pupillometry reflecting the activity of circuits responsible for autonomous stress modulation (68). The PCA prediction maps identify key nuclei coupled with pupil dynamics at different states and also highlight the complexity of brain activation patterns responsible for autonomous and brain state regulation.

Although the present study is based on the anesthetized rat model, it provides a framework that could be applied to analyze human datasets. In particular, the cognitive component of brain activity reflected in pupil diameter changes of awake human subjects could be investigated using the PCA-based fMRI decoding method. Working with awake subjects would additionally mitigate the potential impact of anesthesia on the activity of the sympathetic system (69) which controls pupillary movements in an antagonistic relationship with the parasympathetic system (28, 29). Further research should also be directed towards investigating the state-dependent coupling of pupil dynamics and brain activity at finer temporal scales. Importantly, assuming stationarity of the relationship at any scale could lead to oversimplification of the results, as already evidenced by our ability to differentiate four distinct pupil-fMRI coupling patterns instead of one common correlation map. Combining the analysis of individual fMRI frames (70, 71) with the phase of pupil diameter fluctuation, which is known to reflect the activity of different cortical neural populations (12), would demonstrate whole-brain activity patterns coupled to pupil dilation and constriction. Finally, regions like the subiculum, which previously have not been linked to pupil dynamics, but displayed strong coupling weights in our study, could guide the future electrophysiological studies to reveal novel neuronal regulatory mechanisms underlying pupil dynamics.

## Conclusion

We provided a framework to investigate the brain state-dependent relationship between pupil dynamics and fMRI. The pupil-related brain activity was decoupled from other signal sources based on PCA decomposition and the cluster-specific pupil-fMRI relationship was identified by integrating optimized PCA weighting features using decoding methods. Eventually, distinct subcortical activation patterns were revealed to highlight varied neuromodulatory nuclei corresponding to pupil dynamics.

## Materials and methods

### Animal preparation

All experimental procedures were approved by the Animal Protection Committee of Tuebingen (Regierungsprasidium Tuebingen) and performed following the guidelines. Pupillometry and fMRI data acquired from 10 Sprague Dawley rats had been previously published (23). The rats were imaged under alpha-chloralose anesthesia. For details related to the experimental procedures refer to Pais-Roldan et al. (23).

### fMRI acquisition & preprocessing

All MRI measurements were performed on a 14.1T / 26cm magnet (Magnex, Oxford) with an Avance III console (Bruker, Ettlingen) using an elliptic trans-receiver surface coil (~2×2.7cm). To acquire functional data, a whole-brain 3D EPI sequence was used. The sequence parameters were: 1s TR, 12.5ms TE, 48×48×32 matrix size, 400×400×600μm resolution. Each run had a length of 925 TRs (15 min 25 s). The RARE sequence was used to acquire an anatomical image for each rat. The RARE parameters were: 4s TR, 9ms TE, 128×128 matrix size, 32 slices, 150μm in-plane resolution, 600μm slice thickness, 8× RARE factor. The data from all rats were spatially co-registered. First, for each EPI run, all volumes were registered to the EPI mean. The EPI means were registered to corresponding anatomical images. To register all data to a common template, all RARE images were registered to a selected RARE image. The obtained registration matrices were then applied to the functional data. A temporal filter (0.002, 0.15 Hz) was applied to the co-registered data. The registration was performed using the AFNI software package (72). Principal component analysis (PCA) implemented in Python scikit-learn library (73) was used to reduce the dimensionality of fMRI data for prediction purposes. The PCA time courses were variance normalized before the optimization of linear regression and GRU weights.

### Pupillometry acquisition & pupil diameter extraction

For each fMRI scan, a video with the following parameters was recorded: 24 bits per pixel, 240×352 pixels, 29.97 frames/s, RGB24 format. A customized MRI-compatible camera was used. For details related to the setup refer to Pais-Roldan et al. (23). The DeepLabCut toolbox (74, 75) was used to extract the pupil position from each video frame. The toolbox’s artificial neural network was optimized using 1330 manually labeled images extracted from 74 eye monitoring videos. Training frames were selected using an automated clustering-based DeepLabCut procedure. 4 pupil edge points were manually labeled in each training image. Using the trained network, the 4 points were located in each recorded frame and their coordinates were used to calculate the pupil diameter as:

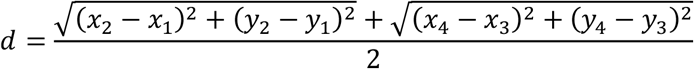

The pupil diameter signals were averaged over 1 s windows to match the fMRI temporal resolution while reducing noise. Pupillometry time courses were variance normalized before the optimization of linear regression and GRU weights.

### UMAP dimensionality reduction

The uniform manifold approximation and projection (UMAP) (44) algorithm was employed to reduce the dimensionality of pupil-fMRI correlation maps before clustering. We used the Python implementation of the algorithm provided by the authors of the method. First, UMAP finds a k-nearest neighbor graph. Based on silhouette scores we set k=5. To facilitate clustering we set the minimum allowed distance between points on the low dimensional manifold to 0. We projected the data from the voxel space (n=20804) to a 72-dimensional representation, as this was the highest number of dimensions the method permitted given 74 input trials.

### Gaussian mixture model clustering

To cluster the trials in the low dimensional space resulting from the UMAP embedding, we used the expectation-maximization algorithm fitting mixture of Gaussians models to the data (45). We used the Python implementation from the scikit-learn library (73) with default parameters.

### Silhouette analysis – cluster number verification

To find the number of clusters for successive analyses we evaluated clustering results using silhouette analysis (46) implemented in the Python scikit-learn library (73). For each point, the method computes a silhouette score which evaluates how similar it is to points in its cluster versus points in other clusters. The clustering of the entire dataset was evaluated by computing the mean silhouette score across all points. The clustering result with the highest mean silhouette score was selected for successive analyses.

### Power spectral density estimation

The spectral analysis was performed using the Python SciPy library (76). To compute the PSDs of utilized signals we employed Welch’s method (77) with the following parameters: 512 discrete Fourier transform points; Hann window; 50% overlap.

### Correlation map-based prediction

Following a strategy described in previous studies (23, 48) we used a pupil-fMRI correlation map to predict pupillometry time courses given fMRI input data. To create the correlation map pupillometry and fMRI data were concatenated across all trials and the pupil diameter fluctuation signal was correlated with each voxel’s signal. This generated a 3D volume (the correlation map) which was then correlated with each individual fMRI volume yielding a single predicted value for each time point. As the resulting time courses’ amplitudes were bounded to the <−1; 1> range and not informative of the target signals amplitudes, Pearson’s correlation coefficient was used to evaluate the quality of the predictions on a trial-by-trial basis.

### Linear regression variants

Linear regression was used to predict pupillometry data given fMRI-PCA inputs. Four linear regression variants were available to a Bayesian optimizer which selected both the linear model type as well as its parameters. The available variants were ordinary least squares, Ridge, Lasso and elastic-net regression models. Python scikit-learn library (73) implementations were used. L2 Ridge regression with a regularization parameter *α* = 19861 obtained the best prediction scores and was found using the Hyperopt toolbox (49, 50).

### GRU

The second model employed for pupillometry decoding was the gated recurrent unit (GRU) (43) artificial neural network. The GRU is a recurrent neural network which encodes each element of the input fMRI-PCA sequence ***x*** into a hidden state vector ***h(t)*** through the following computations:

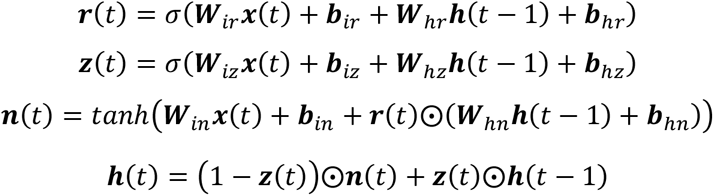

where ***r, z, n*** are the reset, update and new gates, ***W*** are matrices connecting the inputs, gates and hidden states, *σ*() and *tanℎ*() are the sigmoid and hyperbolic tangent functions, ***b*** are bias vectors and ⨀ is the elementwise product. A linear decoder generated predictions based on the hidden state vector:

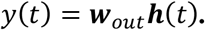

The correlation coefficient was used as the loss function. The networks were trained in PyTorch (78) using the Adam optimizer (79). Hyperparameters were found using Bayesian optimization using the tree of Parzen estimators algorithm (Hyperopt toolbox, n=200) (49, 50). The optimized hyperparameters have been described in Table 1. Early stopping was used in the Bayesian optimization procedure. To set the final number of training epochs for the best network, cross-validation was repeated and the GRU was trained for 100 epochs on each split. Training for 7 epochs yielded the best performance.

**Table 1.** Optimized GRU hyperparameters.

| Parameter name | Description | Range | Final value |
| --- | --- | --- | --- |
| Number of layers | Multiple recurrent layers could be stacked on top of each other. | [1; 3] | 1 |
| Hidden size | Hidden state vector size. | [10; 500] | 300 |
| Learning rate | The rate at which network weights were updated during training. | [10<sup>-6</sup>; 1] | 0.0023 |
| L2 | Strength of the L2 weight regularization. | [0; 10] | 0.0052 |
| Gradient clipping | Gradient clipping (80) limits the gradient magnitude at a specified maximum value. | [yes; no] | yes |
| Max. gradient | Value at which the gradients are clipped. | [0.1, 2] | 1 |
| Dropout | During training, a percentage of units could be set to 0 for regularization purposes (81). | [0; 0.2] | 0 |
| Residual connection | Feeding the input directly to the linear decoder bypassing the RNN's computation. | [yes; no] | no |
| Batch size | The number of training trials fed into the network before each weight update. | [3; 20] | 12 |

### Cross-validation

The available 74 trials were divided into training (n=64) and test (n=10) sets. Linear regression and GRU parameters were found based on the training set with 4-fold cross-validation. The final performance was evaluated on the test set. Scores of the correlation template-based prediction were based on the same data splits.

### Spatial map - linear regression

To create spatial maps highlighting areas that contributed to linear regression predictions we weighted PCA component maps by their associated linear regression weights, summed them and took their means. Region borders from the rat brain atlas (82) were matched to and overlaid on spatial map slices.

### Spatial map – GRU

To create spatial maps highlighting areas that contributed to GRU predictions we computed gradients of each of the predicted time points with respect to the 300 input features. We then averaged the gradients across all time points for each of the features and used these mean values just like the weights in the case of linear regression map generation.

### Variance explained

We obtained the fMRI variance explained by each PCA component directly from the scikit-learn (73) PCA model. To compute the pupil variance explained by each of the PCA time courses we used an approach described in Musall et al. (11) with 4-fold cross-validation. The explained variance of each component was found by randomly shuffling the time points of all other components, training the Ridge linear regression model (*α* = 19861) on shuffled data and assessing the explained variance based on generated predictions.

### Statistical tests – prediction

We used a paired t-test to compare the prediction scores across methods.

### Statistical tests – linear regression spatial maps

To test which linear regression spatial map values significantly contributed to the predictions we used randomization tests. For each cluster, we shuffled the input and output pairings 10000 times, trained a linear model and created a spatial map for each of those pairings. We then compared the values in the original maps with the shuffled ones. Values that were at least as extreme as the shuffled values at the 0.005 positive or negative percentile (p=0.01) were considered significant. The results were controlled for false discovery rate with adjustment (83, 84).

## Acknowledgments

We thank Dr. R. Pohmann and Dr. K. Buckenmaier for technical support; Dr. E. Weiler, Dr. P. Douay, Mrs. R. König, Ms. S. Fischer, Ms. H. Schulz and Dr. Jörn Engelmann, for animal/lab maintenance and support; the Analysis of Functional NeuroImages (AFNI) team for software support.

## Author contributions

Research design: X.Y., F.S., Data acquisition: P.P., K.T., Analysis: F.S., Writing – original draft: F.S., X.Y., Writing – review & editing: X.Y., F.S., P.P., K.T., Supervision: X.Y.

---

**Supplementary Fig. 1.**
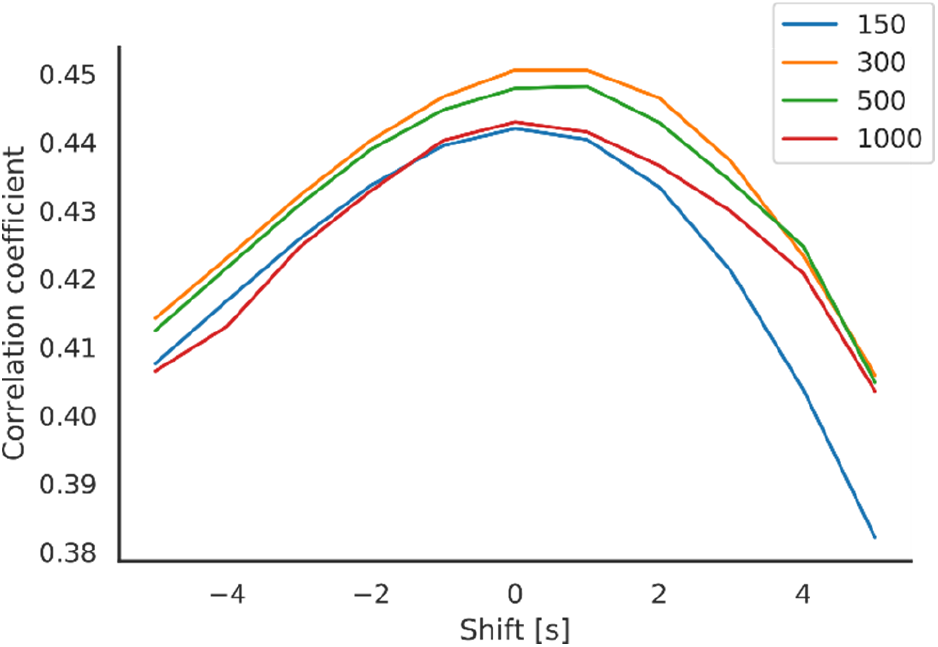
The influence of temporal shifts and different component counts on prediction scores. Temporally shifting the fMRI and pupil fluctuation signals reduces prediction accuracy. Predictions generated based on 300 PCA components obtain the best scores.

**Supplementary Fig. 2.**
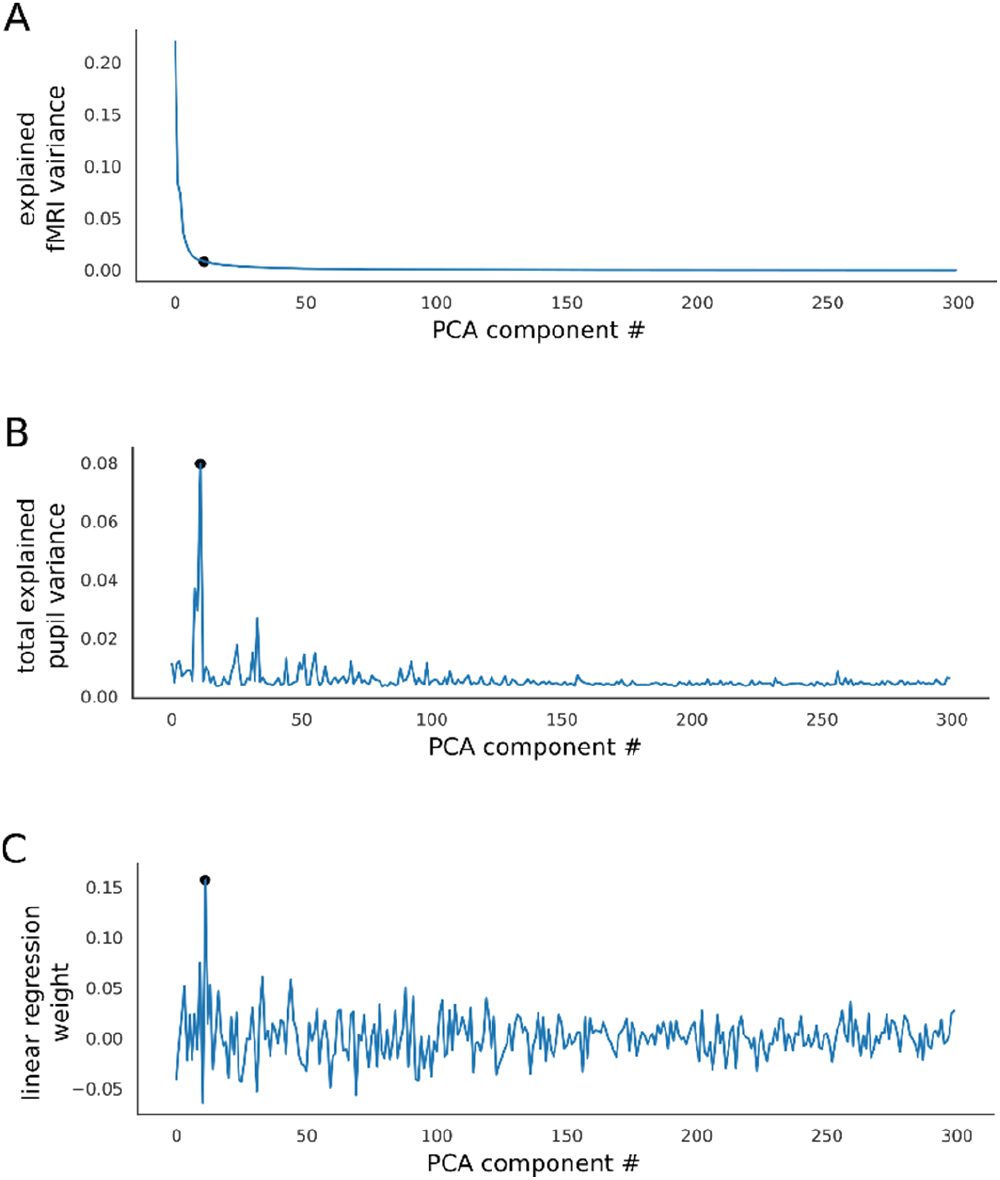
PCA decoupling of pupil-related fMRI activity from other signal sources. (A) fMRI variance explained by individual PCA components. Components are ordered by variance explained. Black dot marks the component with the most explained pupil variance. (B) Pupil variance explained by the PCA signals. Same ordering as in A. The component explaining the most pupil variance (explained var.=7.03 %) explains only 0.8% of fMRI variance. Black dot marks the component with the most explained pupil variance. (C) Weights of the linear regression model trained to decode pupil signals based on fMRI-PCA data. Each weight corresponds to a single PCA component’s time course. The highest absolute weights are not assigned to the components explaining the most fMRI variance. Black dot marks the component with the most explained pupil variance.

**Supplementary Fig. 3.**
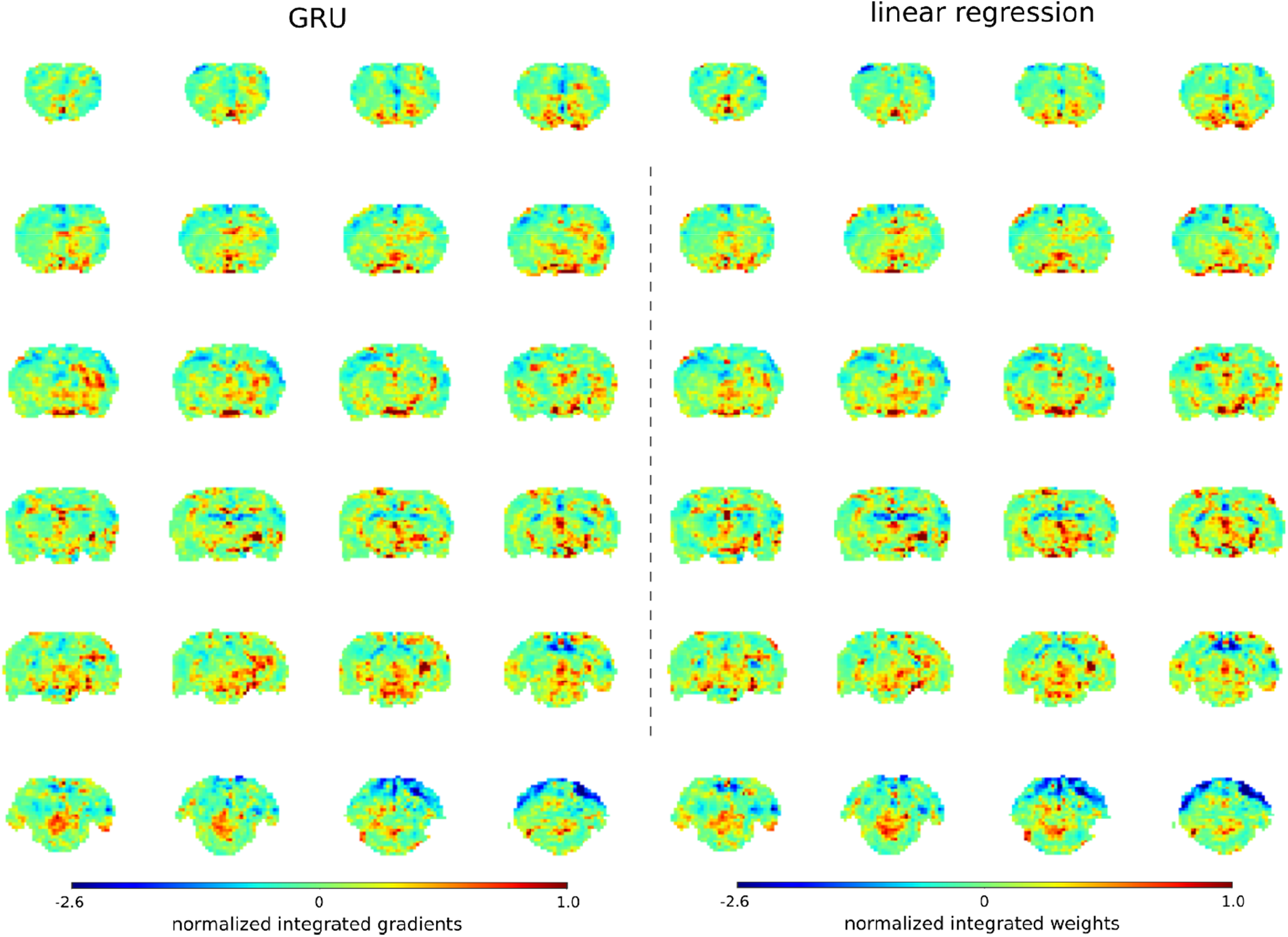
Similarity of GRU and linear regression prediction maps. Spatial maps were generated by integrating PCA spatial maps with either linear regression weights or average GRU gradients. The maps highlight the same areas. This observation coupled with the similarity of predictions generated by both methods and the short GRU training time suggests that a linear mapping was sufficient for this decoding task.

